# Plasmids encode niche-specific traits in *Lactobacillaceae*

**DOI:** 10.1101/2020.08.20.258673

**Authors:** Dimple Davray, Dipti Deo, Ram Kulkarni

**Author notes:** **Corresponding Author Dr. Ram Kulkarni**, or.

## Abstract

The species of family *Lactobacillaceae* are found in highly diverse environments and play an important role in fermented foods and probiotic products. Many of these species have been individually reported to harbor plasmids that encode important genes. In this study, we performed comparative genomic analysis of the publically available data of 512 plasmids from 282 strains represented by 51 species of this family and correlated the genomic features of plasmids with the ecological niches in which these species are found. Two-third of the species had at least one plasmid-harboring strain. Plasmid abundance and GC content were significantly lower in the vertebrate-adapted species as compared to the nomadic and free-living species. Hierarchical clustering (HCL) highlighted the distinct nature of plasmids from the nomadic and free-living species than those from the vertebrate-adapted species. EggNOG assisted functional annotation revealed that genes associated with transposition, conjugation, DNA repair and recombination, exopolysaccharide production, metal ion transport, toxin-antitoxin system, and stress tolerance were significantly enriched on the plasmids of the nomadic and in some cases nomadic and free-living species. On the other hand, genes related to anaerobic metabolism, ABC transporters, and major facilitator superfamily were found to be overrepresented on the plasmids of the vertebrate-adapted species. These genomic signatures are correlated to the comparatively nutrient-depleted, stressful and dynamic environments of nomadic and free-living species and nutrient-rich and anaerobic environments of the vertebrate-adapted species. Thus, these results indicate the contribution of the plasmids in the adaptation of lactobacilli to the respective habitats. This study also underlines the potential application of these plasmids in improving the technological and probiotic properties of lactic acid bacteria.

**Impact statement:** The bacteria of the family *Lactobacillaceae* are present in the wide range of habitats and play an important role in human health, fermented foods and chemical industries. A few studies have demonstrated the presence of plasmids in the individual strains of *Lactobacillaceae* species encoding various traits. Extensive data of genome sequences of the lactobacilli are becoming available; however, no comprehensive analysis of the plasmid-encoded genes and determining their biological relevance across lactobacilli has been undertaken at a larger scale. In this study, we explored the genomic content of 512 plasmids of *Lactobacillaceae* species and correlated it to the three types of these species according to their ecological niches – vertebrate-adapted, free-living and nomadic. Comparatively lower plasmid abundance and GC content in the vertebrate-adapted species could be correlated to the presence of these species in the nutrient-rich environment. The genomic content of the plasmids was consistent with the respective lifestyle adopted by lactobacilli suggesting that the plasmids might enhance the niche-specific fitness of the strains. The plethora of important genes present on the plasmids can also make them a highly useful tool in improving the probiotic, technological and food-related properties of lactobacilli.

**Data summary:** Nucleotide sequences of plasmids of *Lactobacillus* strains for which complete genome sequences were available were retrieved from the NCBI genome [https://www.ncbi.nlm.nih.gov/genome] and PATRIC 3.5.41 databases on 31st March 2019. The dataset includes 512 nucleotide sequences of plasmids of 282 strains belonging to genus *Lactobacillus* before its reclassification into several genera (1). Details of the plasmids have been given in Table S1.

## 1. Introduction

Lactobacilli are gram-positive, aerotolerant, and non-sporulating bacteria which belong to the lactic acid bacteria (LAB) group wherein lactic acid is the major metabolic end product during glucose fermentation (2). *Lactobacillus* is a major genus of the *Lactobacillaceae* family and the highly diverse nature of its species has recently resulted in reclassification of this genus into 23 genera (1). Many lactobacilli have been associated with humans in a variety of ways. Their presence in the gastrointestinal tract has earned extraordinary attention due to the health-promoting properties of few of the genera. These bacteria are also important in the food industry for the production of fermented dairy products wherein these bacteria provide the taste, texture, and antibacterial activity (3). Furthermore, lactobacilli have applications in the production of industrially important chemicals such as lactic acid (4).

Lactobacilli have been isolated from a wide range of habitats and have been found to have highly diverse traits. Based on the properties such as phylogenetic analysis, source of isolation, the prevalence of detection, optimal growth temperature, substrate utilization, and environmental stress tolerance, the species of the earlier *Lactobacillus* genus have been classified as host-adapted, nomadic and free-living (5). Some of the genomic features of these species are correlated with these habitats. For example, the vertebrate-adapted lactobacilli (VAL) have undergone genome reduction by losing a substantial number of genes involved in carbohydrate metabolism and amino acid and cofactor biosynthesis because they thrive in a nutrient-rich environment (5), (6), (7). On the other hand, free-living lactobacilli (FLL) and nomadic lactobacilli (NL) have larger genome sizes because they encode various enzymes which utilize a broad spectrum of carbohydrates (8). Furthermore, *Lactobacillus helvticus* DPC4571 and *Lactobacillus acidophilus* NCFM were found to have genes important for the adaptation to their habitats, *viz*., dairy and gut, respectively (9). Lastly, *Lactiplantibacillus plantarum*, a hallmark NL species found in most diverse habitats was also observed to have a highly diverse genome with no unique environmental signature (10). Another classification of lactobacilli is based on the carbohydrate fermentation pathways, as homofermentative and heterofermentative. In the homofermentative species, the phosphotransferase system (PTS) is preferred to transport the sugars which are fermented via the Emden-Meyerhoff pathway. On the other hand, in the heterofermentative species, PTS are not functional and the phosphoketolase pathway is the preferred catabolic pathway (11), (12).

Plasmids have been extensively found in lactobacilli and reported to carry the genes that give them favorable traits and permit them to survive in competitive surroundings (13). Many plasmids have been sequenced from the individual strains and found to encode important functions such as oxidative stress response, antibiotic resistance, bacteriophage resistance, chloride and potassium transport, and bacteriocin production (14), (15). Only a handful of reports are available on the importance of plasmids in the niche adaptation of lactobacilli. In *L. plantarum* P-8 isolated from fermented dairy products, loss of the plasmids important for dairy-adaptation was found when the strain was administered in rats and humans (16). A plasmid from *Lacticaseibacillus paracasei* NFBC338 isolated from the human gut encoded a gene involved in adherence to the human intestinal epithelial cells (17). In *Levilactobacillus brevis*, the brewery isolates were found to have the unique genes on the plasmids supporting the growth under the harsh conditions. At the same time, a large proportion of brewery plasmidome was also found to be shared with the insect isolates (18). Similarly, in *Lactococcus lactis*, another important bacterium of the LAB group, plant cell wall modifying enzymes were encoded by the plasmids in the plant-derived isolates (19). Apart from these scarce strain and species-specific reports, the contribution of plasmids in niche-adaptation has not been holistically studied in the *Lactobacillaceae* family. In the last decade, Next Generation Sequencing (NGS) has created extensive data on the genome sequences of many strains of lactobacilli. In the current study, we report an analysis of the plasmid sequence data of the species of genus defined as *Lactobacillus* before its division into several genera (1), available in the public domains to explore their genomic content and their possible involvement in the niche adaptation.

## 2. Material and Methods

### 2.1 General characterization of the plasmids

The coding sequences (CDSs) of the 512 plasmids were extracted from the GenBank database, along with their NCBI annotation (https://www.ncbi.nlm.nih.gov/nucleotide) (20). The plasmids for which the encoded genes were not annotated in NCBI were subjected to the analysis using Prodigal software (version 2.6) for predicting the CDSs.

### 2.2 Functional annotation

The functional annotation and classification of the CDSs were performed using eggNOG 4.5 databases (http://eggnogdb.embl.de/). For subgrouping of the genes under each COG category, their annotations in eggNOG4.5, NCBI and KEGG databases were considered.

### 2.3 Markov clustering analysis

The all-against-all bi-directional blast alignment was performed on the obtained CDSs with at least 50% amino acid identity and 50% query coverage (20). The blast output was analyzed using markov clustering (MCL) in the mclblastine v12-0678 pipeline to classify proteins into families (21). Further, Hierarchical clustering was computed in TM4 MeV Suite (22) based on the presence and absence of the protein families in plasmids by using average linkage algorithm and Manhattan distance metric parameters The HCL analysis was visualized in the Interactive Tree of life (ITOL) (23) by importing Newick tree from TM4 MeV Suite. For identification of the plasmids which are possibly shared between the species from different habitats, a similarity matrix was created for the plasmids based on the presence and absence of the MCL families using the Sørensen–Dice coefficient. The Genbank files of the plasmids with Sørensen–Dice similarity coefficient of 0.8 or more were compared to each other using EasyFig program (24).

### 2.4 Identification of other important genes

To identify antibiotic resistance gene, initially, amino acid sequences of 2,404 antibiotic resistance (ARGs) genes were downloaded from the CARD database (http://arpcard.mcmaster.ca). Plasmids CDSs were aligned against retrieved ARGs sequences through standalone blast (BLASTP,ver.2.9.0, https://ftp.ncbi.nlm.nih.gov/blast/executables/LATEST/) with a threshold of query coverage of >70%, amino acid sequence identity of >30%, and e-value <10^−5^ The exopolysaccharide (EPS) gene clusters were identified as described earlier (25). Bacteriocin operons were identified using the web version of BAGEL4 (http://bagel4.molgenrug.nl) using default parameters.

### 2.5 Statistical analysis

GraphPad Prism 8 was used to perform statistical analysis.All the comparisons between the habitats was performed by Kruskal-Wallis test with Dunn’s post hoc test. Wilcoxon Signed Rank Test was used to compare GC content of the plasmids with that of the chromosome of the same strain and species.

## 3. Result and Discussion

### 3.1 Data collection and general characterization

Till March 2019, NCBI prokaryotic genome database was found to have complete genome sequences for 282 strains under the genus defined as *Lactobacillus* before its reclassification into several genera (1). With the current classification, this data includes 24 of the 31 genera under the *Lactobacillaceae* family. Collectively, nucleotide sequences of a total of 512 plasmids were found in the database for all these strains (Table 1, Table S1). Based on the classification by Duar et al. (5), in our dataset, the maximum number of strains(110) belonged to NL group followed by VAL (86),FLL (43) and insect adapted (6) groups. Forty strains belonged to the species for which no clear information on their possible native habitat was available; hence they were not considered for further analysis. Plasmids from insect adapted strains were excluded from the subsequent analyses as only three plasmids were represented by this category of the strains. Similarly, based on the glycolytic pathway, 216 strains belonged to homofermentative and 51 to heterofermentative species (11), (12).

**Table 1:**
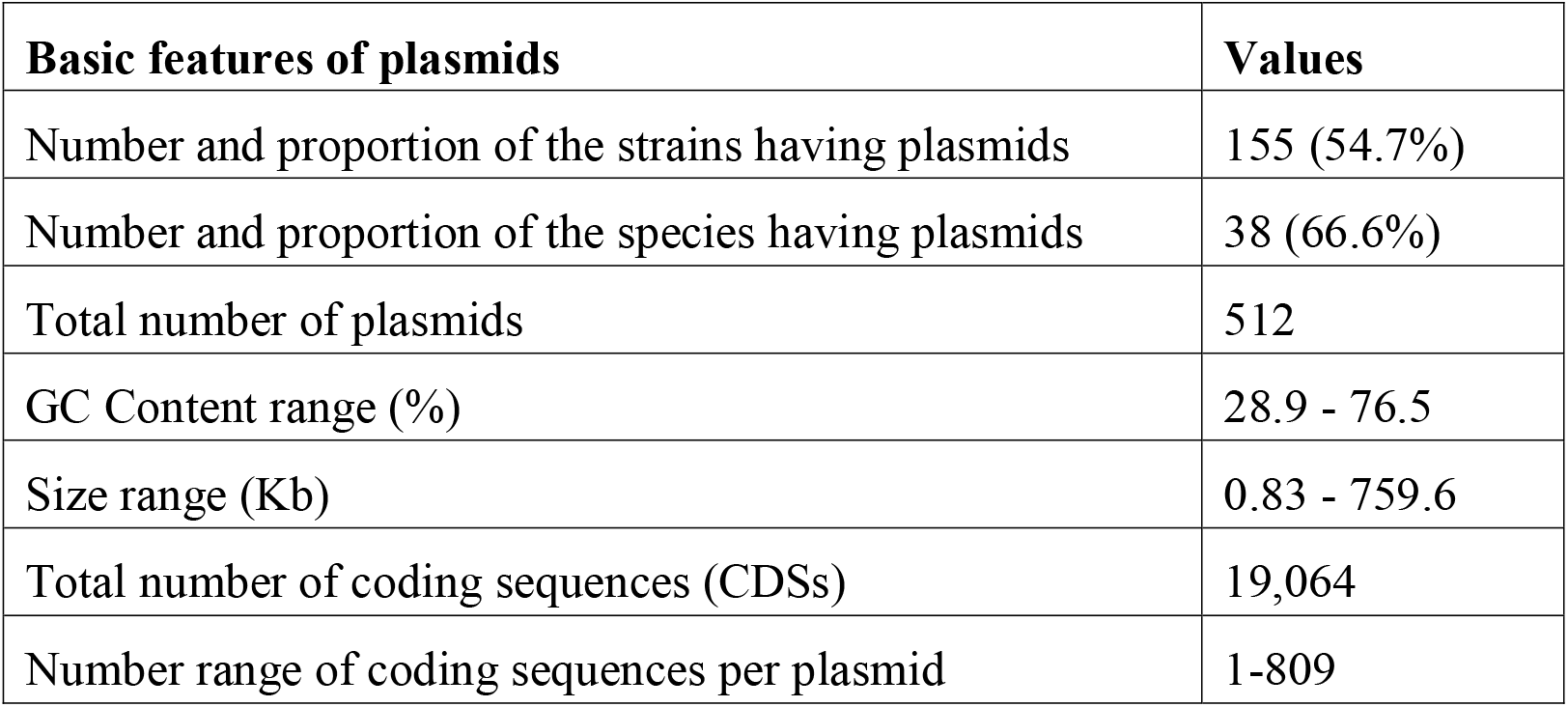
General features of the plasmids from completely sequenced *Lactobacillaceae* species.

| Basic features of plasmids | Values |
| --- | --- |
| Number and proportion of the strains having plasmids | 155 (54.7%) |
| Number and proportion of the species having plasmids | 38 (66.6%) |
| Total number of plasmids | 512 |
| GC Content range (%) | 28.9 - 76.5 |
| Size range (Kb) | 0.83 - 759.6 |
| Total number of coding sequences (CDSs) | 19,064 |
| Number range of coding sequences per plasmid | 1-809 |

The proportion of strains having plasmids was the smallest for VAL (32.2%) and the largest for FLL (76.7%) (Fig. 1a). The average number of plasmids per strain was significantly lower for VAL (1) than NL (2.3) and FLL (2.7) (Fig. 1b, Table S1). On the other hand, the average number of plasmids per strain for each species was significantly higher only in FLL as compared to VAL, though this average was three-fold higher for NL than VAL (Fig. 1c). This suggests that some of these differences could be driven by strains of only a few species; whereas some others are possibly common to majority of the species of that habitat. Overall, the lower plasmid abundance in VAL as compared to NL and FLL is in agreement with the similar observation for the chromosomal genome size (5). Such reduction is considered to be due to the loss of several biosynthetic pathway genes as the bacteria are living in a nutrient-rich environment in the hosts. Furthermore, loss of plasmids in a dairy isolate of *Lactobacillus* when administered in the human and rat has also been reported. Such loss has been postulated to be because of the superfluous nature of the genes encoded by the lost plasmid in the gut environment as well as stress response of the bacteria during the passage through the harsh gastric environment (16). There was no difference in the average plasmid size per strain across various habitats (Fig. 1d), suggesting that the low number of plasmids in VAL is not compensated by their larger size in these strains.

**Figure 1:**
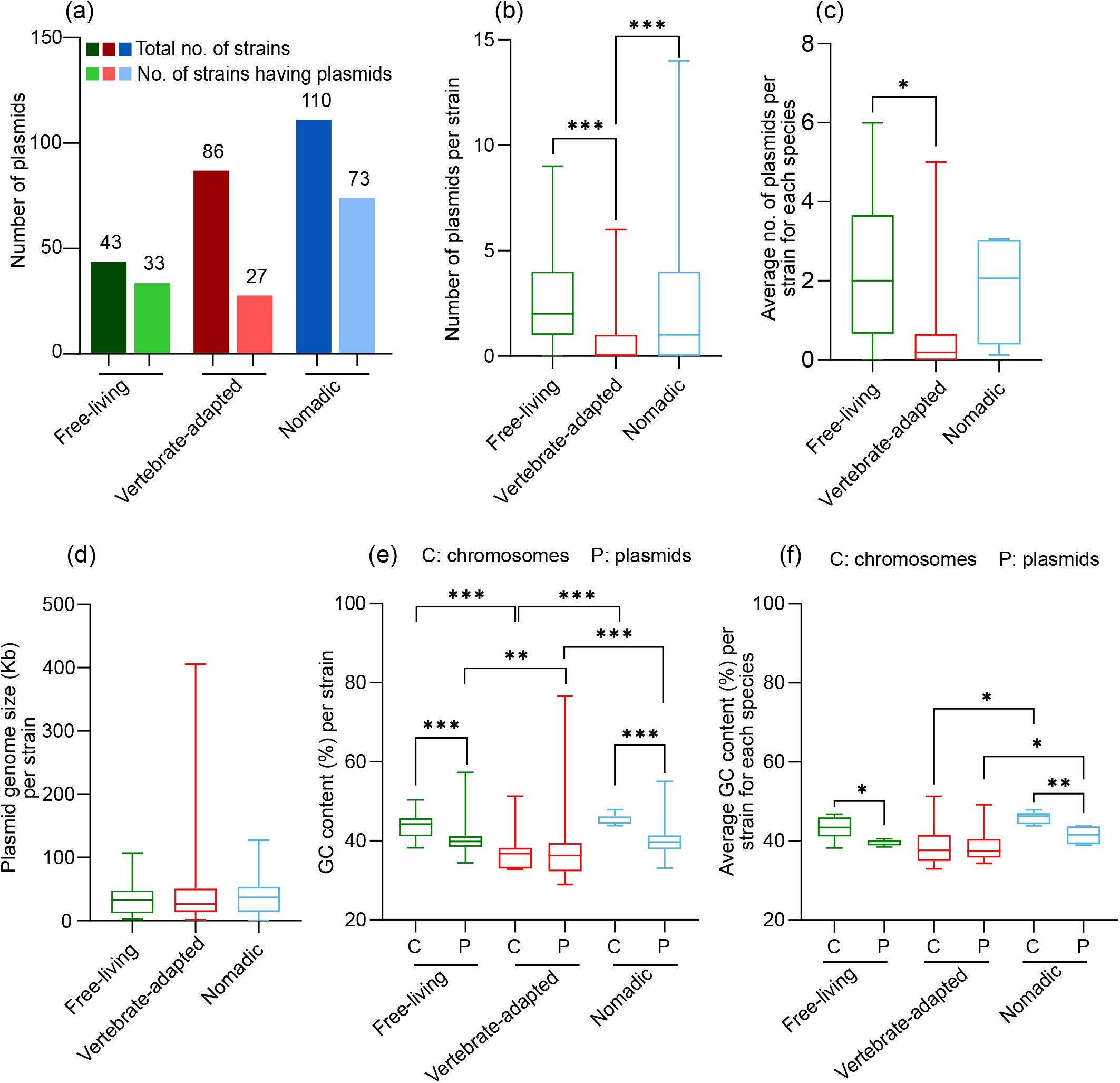
General features of the plasmids in the *Lactobacillaceae* species from various habitats indicating the abundance of plasmids across habitats (a), the number of plasmids per strain (b), average number of plasmids per strain for each species (c), average size of the plasmids per strain (d), GC content per strain (e) and average GC content per strain for each species (f). The statistical analyses were performed to compare the data across various habitats using Kruskal-Wallis test with Dunn’s post hoc test. The GC contents of plasmids and chromosome of the same strain (e) and of the same species (f) were compared using the Wilcoxon signed rank test (***, p < 0.001; **, p < 0.01; *, p < 0.05).

The average GC content of the plasmids per strains was significantly lower for VAL (37.7%) than NL (39.7%) and FLL (39.7 %) (Fig. 1e). At the species level, this difference was significant only between VAL and NL (Fig. 1f). This observation is similar with the lower chromosomal GC content of the host-associated strains, which is thought to be because of non-adaptive loss of DNA repair system leading to the mutational bias toward A and T (5). The plasmid GC contents per strain and per species for NL and FLL were lower than the respective chromosomal GC contents; whereas, in VAL, plasmids and chromosomes had similar GC content (Fig. 1e and 1f). This observation was also valid for similar comparison within the individual strains wherein the average plasmids GC content was found lower than the respective chromosomal GC content in all the NL and in 27 of 33 FLL. It has been reported that the maintenance of plasmid with higher GC content is metabolically expensive for the bacterial cells with lower chromosomal GC content and is thus evolutionary unfavourable (26). It has also been proposed that plasmids can be transferred from chromosomally GC-poor hosts or environments to relatively GC-rich hosts (27). Thus, the lower GC content of plasmids from NL and FLL than their chromosomes is not surprising and is indeed an indication of their possible acquisition by horizontal gene transfer. On the other hand, it is suggested that with time the nucleotide composition of plasmids might become closer to that of the host genome because of the dependency of plasmid replication on the host cell machinery (28), (29). Thus, the comparable GC content of VAL chromosomes and plasmids probably suggests their long association with each other. No correlation was observed between GC content and size of the plasmids (data not shown) in contrast to the earlier observation (30).

### 3.2 Hierarchical clustering (HCL) analysis

To understand the extent of similarity between the plasmids based on the genomic content, their functional grouping was performed by HCL analysis. Based on markov clustering analysis (MCL), the coding sequences (total 19,058) could be classified into a total of 3,380 protein families, with about half of them (1,560) having a single-member. This observation is similar to that on *Lc. lactis* plasmids, where also approximately half (413 out of 885) of the protein families were singleton (31). Plasmids from VAL had the highest proportion (73%) of unique families, followed by NL (55%) and FLL (44%) (Fig. 2a). This probably suggests the uniqueness of the plasmids in VAL, which might have arisen during the long course of association of plasmids with these species as speculated from the GC content. The highest sharing of families of NL with VAL and FLL could be due to the fact that NL is found in a wide range of environments, including those associated with the hosts as well as those similar to FLL (5).

**Figure 2:**
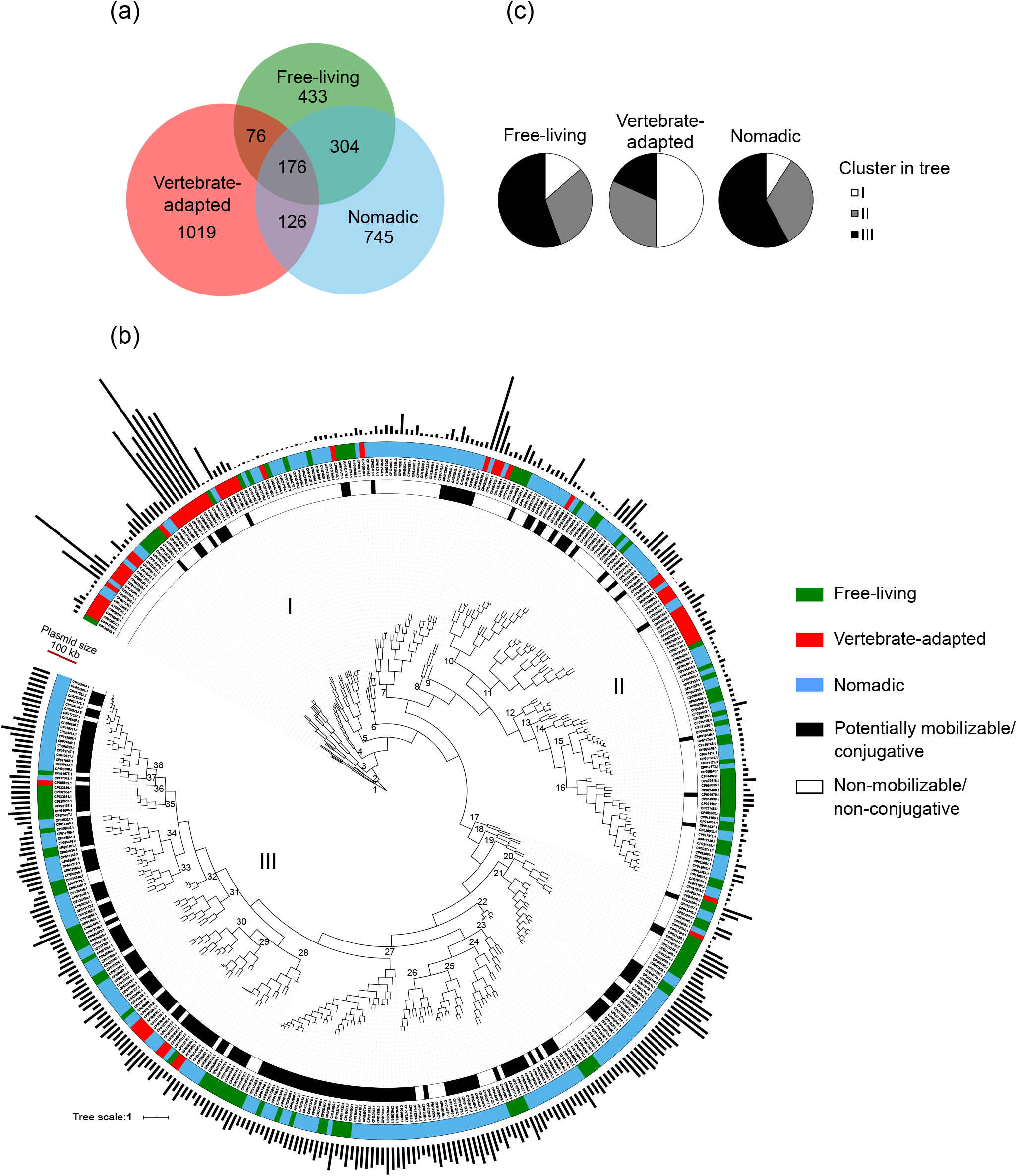
Hierarchical clustering (HCL) analysis of the *Lactobacillaceae* plasmids. (a) Sharing of protein families encoded by plasmids as identified by markov clustering (MCL) across lactobacilli from various habitats. (b) HCL tree displaying the clustering of the plasmids based on their protein family composition. Black bars on the outermost periphery represent plasmid genome size. The plasmids having mobilization/conjugation genes were identified based the eggNOG annotations detailed in section 2.2.1 (c) Pie chart representing the proportion of the plasmids from each habitat under clusters I, II, and III of the tree shown in panel (b).

The data obtained on protein families from MCL analysis was used to perform HCL to cluster the plasmids based on the presence and absence of protein families (Fig. S1). The analysis depicted three broad clusters (I, II and III) of the plasmids (Fig. 2b). Although there was no strict distinct clustering based on the habitats, certain habitat-wise observations were made. For example, half of the plasmids from VAL were present in the cluster I whereas the majority of the plasmids from NL and FLL (57.9 % and 55.5 %, respectively) were present in cluster III. On the other hand, almost equal proportions of plasmids from VAL (31.7 %), NL (33.1%) and FLL (31.1 %) were present in cluster II (Fig. 2c). Within the three main clusters, 38 subclusters were observed (Fig. 2b, Table S2). Out of 100 strains having two or more plasmids, 63 strains were such that none of the plasmids from the same strain belonged to the same subcluster. This observation probably indicates the non-redundant genomic content of multiple plasmids within a strain in the majority of lactobacilli. Similar observation was reported in *Lc. lactis* IL594 where multiple plasmids in the same strain were found to be diverse and proposed to function in the synergistic manner (32).

Based on the Sørensen–Dice similarity coefficient which was a measure of extent of sharing of the protein families between plasmids, 15 plasmid pairs with the Sørensen–Dice coefficient of higher than 0.8 were identified (Table S3). Of all these plasmid pairs, four having Sørensen–Dice coefficient of one had only one protein family and the sequence identity of < 85% with the partner plasmid with query coverage of < 60%, hence they were considered dissimilar. Majority of the remaining pairs (6) had plasmids from the species belonging to dissimilar habitats. BLAST analysis of the plasmids found to be similar to each other in this way using EasyFig program suggested their highly similar gene content (Fig. S2). This is a suggestive of the inter-species as well as inter-habitat sharing of some of the plasmids in lactobacilli. Interestingly, none of the plasmids from these pairs belonged to VAL underlining the uniqueness plasmids from VAL. This further supports the evolutionary model proposing that VAL have evolved independently in the confined vertebrate-associated environment (5).

### 3.3 Functional annotation

To predict the putative functions associated with the plasmid-encoded genes, they were classified based on orthology using eggNOG. Based on this, 63.2% (12,120) of CDSs were annotated into 20 Clusters of Orthologous Groups (COGs) functional categories. Replication, recombination, and repair was the largest category with 32.5% of the CDSs followed by Function unknown (20.7% CDSs), Carbohydrate transport and metabolism (6.2% CDSs) and Transcription (5.3% CDSs) (Fig. 3, Table S4). Furthermore, the genes were subgrouped under each eggNOG category based on their putative functions based on eggNOG, NCBI, and KEGG annotations. The genes under Function unknown category were manually evaluated for their annotations in these databases and based on this some of the genes were binned into the individual COG categories. For example, genes annotated as LPXTG-motif cell wall anchor domain protein that were found under the Function Unknown category were manually put under Cell wall/membrane/envelope biogenesis.

**Figure 3:**
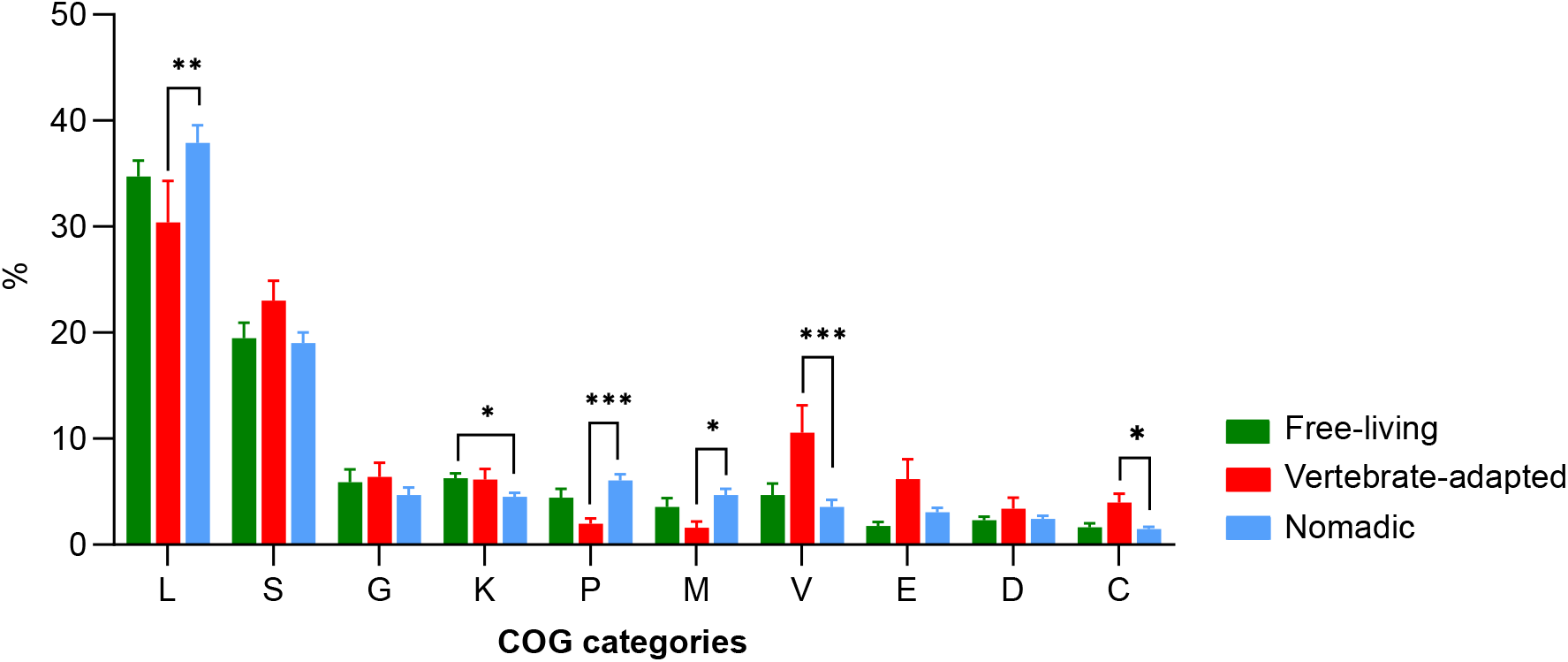
Average proportion of each of the top-ten COG categories in the *Lactobacillaceae* plasmids per strain from various habitats. L, Replication, repair and recombination; S, Function unknown; G, Carbohydrate transport and metabolism; K, Transcription; P, Inorganic ion transport and metabolism; M, Cellwall/membrane/envelope biogenesis; V, Defense mechanisms; E, Amino acid transport and metabolism; D, Cell cycle control, cell division, chromosome partitioning; and C, Energy production and conservation. The statistical analysis was performed to compare the data across various habitats using the Kruskal-Wallis test with Dunn’s post hoc test (***, p < 0.001; **, p < 0.01; *, p < 0.05). The error bars represent standard error of measurement.

To test the hypothesis that plasmids play an important role in the environmental adaptations in *Lactobacillaceae*, we investigated whether the plasmids encode genes with varying putative functions across the lifestyles. The fraction of genes involved in Replication, recombination, and repair (L), Cell wall/membrane/envelope biogenesis (M), and Inorganic ion transport and metabolism (P) was significantly higher (p < 0.05) in the plasmids from NL as compared to VAL (Fig. 3). Furthermore, the genes associated with Defense mechanisms (V) and Energy production and conservation (C) were significantly overrepresented (p < 0.001, and p < 0.05 respectively) in VAL in comparison with NL (Fig. 3). Similarly, the average proportion of genes involved in Transcription (K) was significantly higher (p < 0.05) in the plasmids from FLL as compared to NL.

#### 3.3.1 Replication, recombination, and repair

Various subcategories under Replication, recombination, and repair (L) were transposase (1,928 CDSs), DNA replication (614 CDSs), resolvase (323 CDSs), integrase (256 CDSs), mobilization/conjugation (221 CDSs), DNA repair (191 CDSs) and DNA repair and recombination (99 CDSs) (Fig. 4a, Table S5). Within these, the genes encoding transposases, resolvases, DNA repair and recombination proteins, and mobilization/conjugation proteins were significantly overrepresented on the plasmids from NL than those from VAL considering the strain-level average. Of these genes, resolvase and transposase were significantly overrepresented on the plasmids from FLL as compared to VAL at species level also (Fig. 4a, Table S6). Previous studies have demonstrated that transposases are more abundant in the bacteria living in the extreme environment and they can help in the evolution of fitter phenotypes by accelerating the rate of mutations (33), (34). In the context of lactobacilli, NL and FLL can be considered relatively extreme as these species occur in a wider range of harsh environmental conditions (5). Thus, higher proportion of mobile genetic elements in the plasmids of NL and FLL may give them genomic plasticity for adaptation to the dynamic environments. Since NL and FLL live are under more extreme conditions, they are likely to face more DNA damage, which can justify the higher proportion of DNA repair genes in their plasmids. This speculation is also supported by earlier observations reporting a higher proportion of DNA repair genes in the thermophilic microorganisms (35). Furthermore, the low proportion of DNA repair genes in VAL could also be correlated to the reduced genome size of these bacteria. The correlation between smaller proteome and loss of the DNA repair genes has already been established in several other bacteria (36).

**Figure 4:**
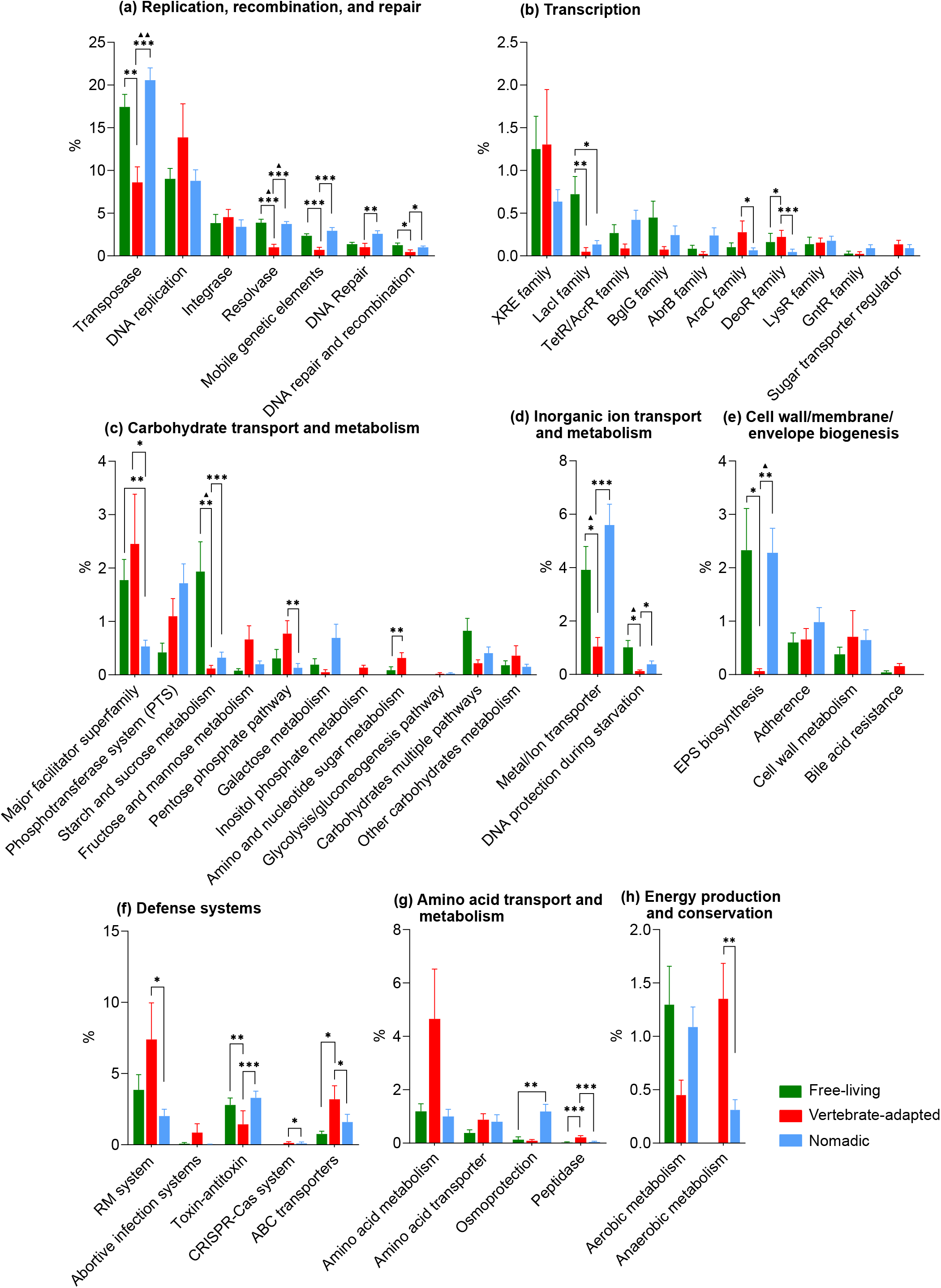
Average proportion of each of the COG subcategories in the *Lactobacillaceae* plasmids per strain from various habitats. The statistical analyses were performed to compare the data across various habitat using the Kruskal-Wallis test with Dunn’s post hoc test (***, p < 0.001; **, p < 0.01; * p < 0.05). The error bars represent standard error of measurement. The solid black triangles indicate the significant difference by the same test between the habitats for the species-level average which is mentioned in the Table S6 (^▲▲▲^, p < 0.001; ^▲▲^, p < 0.01; ^▲^, p < 0.05).

#### 3.3.2 Carbohydrate transport and metabolism

In the current study total of 673 genes (6.2% of the total plasmid-encoded genes) associated with Carbohydrate transport and metabolism (G) were found. Further, carbohydrates metabolism (332 CDSs), phosphotransferase system (PTS) (177 CDSs) and major facilitator superfamily transporter (131 CDSs) were identified as the major subcategories (Fig. 4c, Table S5). Fourty-nine complete PTS transporters were found in the 35 plasmids across 30 strains (Table S7). Although there was no significant difference between the habitats for the average proportion of PTS genes, this proportion per species was significantly higher for the NL as compared to HAL and FLL. This observation was also reflected in the fact that the proportion of species having complete PTS on plasmids was 22.2%, 9.1%, and 83.3% for FLL, HAL, and NL, respectively (Table S7). NL are metabolically versatile and able to utilize a wide range of sugars. Thus, the broader distribution of complete PTS on the plasmids from NL underlines the contribution of plasmids in adaptation to diverse environments. All the 30 strains having complete PTS were homofermentative which is consistent with the reported loss of PTS genes from the chromosomes of heterofermentative species (11). These results suggest that plasmids can significantly contribute to the physiology of the homofermentative species by allowing sugar uptake via PTS systems.

The abundance of the genes from the KEGG category ‘starch and sucrose metabolism’ in *L. brevis, L. sakei, Secundilactobacillus paracollinoides, L. paracasei*, which are FLL can be clearly attributed to the presence of maltose phosphorylase and β-phospho-glucomutase in their plasmids which allow utilization of maltose. Furthermore, a large number of maltose/moltooligosaccharide transporters were encoded on the plasmids of FLL (Table S5). Plasmids having such maltose utilization genes had *lacI* on their upstream region in the six strains of *L. sakei* which is one of the FLL (Fig. 4b, Table S8). LacI transcriptional regulator was shown to be involved in the maltose utilization in *L. acidophilus*, *Enterococcus*, and *Bifidobacterium* (37). One of the major habitats for FLL is fermented cereals (5), which are likely to be rich in maltose released by starch hydrolysis. This suggests that the plasmids might contribute to the efficient utilization of maltose in FLL.

The complete lactose operon (*lacTEFG*) was found on the plasmids of six strains of *L. paracasei* and one strain of *Lacticaseibacillus rhamnosus* (Table S9). A previous study has shown that this operon also helps *Lacticaseibacillus casei* in the uptake of N-acetyllactosamine in human milk (38). AraC was present upstream of the rhamnose utilization genes, *viz*., MFS transporter, rhamnulokinase, L-rhamnose mutarotase, L-rhamnose isomerase, rhamnulose-1-phosphate aldolase in five plasmids from *Ligilactobacillus salivarius* (Fig. 4b, Table S10). AraC transcriptional regulator was shown to be involved in the rhamnose metabolism in *L. acidophilus* and *L. plantarum* (39). These results suggest that plasmids might play an important role in the growth and physiology of *Lactobacillaceae* species by allowing the uptake of various sugars.

Genes encoding major facilitator superfamily (MFS) proteins were found in significantly higher proportion on the plasmids of VAL as compared to NL and FLL (Fig. 4c). MFS proteins are involved in the transport of a large number of molecules across the membrane in response to chemiosmotic gradient. These include sugars, vitamins, antibiotics, amino acids, nucleosides, ions, etc. In general, majority of the MFS transporters in LAB are multidrug transporters (40). Thus, their overrepresentation in the plasmids of VAL might be related to the antibiotic usage in humans and other animals. Additionally, some of the MFS transporters are also involved in the bile salt resistance in *Lactobacillus* (41) which is relevant for VAL.

#### 3.3.3 Inorganic ion transport and metabolism

Within the Inorganic ion transport and metabolism (P) category metal/ion transporters (435 CDSs), other transporters (87 CDSs) and DNA protection during starvation protein (*dps*) (46 CDSs) were identified as the dominant genes (Table S5). Metal/ion transporters genes were significantly more abundant in the plasmids from NL and FLL than VAL considering strains-level average (Fig. 4d, Table S6). This difference was also reflected in species-level average between FLL and VAL. Under this category, a large number of genes putatively involved in heavy metal resistance (HMRGs) were found including those ascribing resistance to arsenic (98 genes), cadmium (49 genes), copper (19 genes) and cobalt (13 genes) on the plasmids of 69 strains (Table S5). Such heavy metal resistance genes, especially those involved in arsenic resistance were highly enriched on the plasmids of NL such as *L. plantarum, L. paracasei, L. paraplantarum* as compared to VAL (65 and 4 genes, respectively). No comparative phenotypic data on the heavy metal resistance of the *Lactobacillaceae* species belonging to various habitats is available. However, the above observation is correlated to a finding in the case of *Acinetobacter* and *Ornithilibacillus* where in the environmental isolates were more resistant to heavy metals than the clinical isolates (42), (43). Further, the gene clusters involved in arsenate resistance were detected in 15 plasmids from NAL and FLL (Fig. 5, Table S11). Eight plasmids had complete clusters encoding all five proteins, *viz*., a regulator (*arsR*), ATPase component (*arsA*), secondary transporter (*arsB*), transcriptional regulator (*arsD*), and arsenate reductase (*arsC*). The other seven plasmids lacked *arsC*; however, chromosomal *arsC* can complement such incomplete cluster as shown earlier in the case of *L. plantarum* WCFS1 (44).

**Figure 5:**
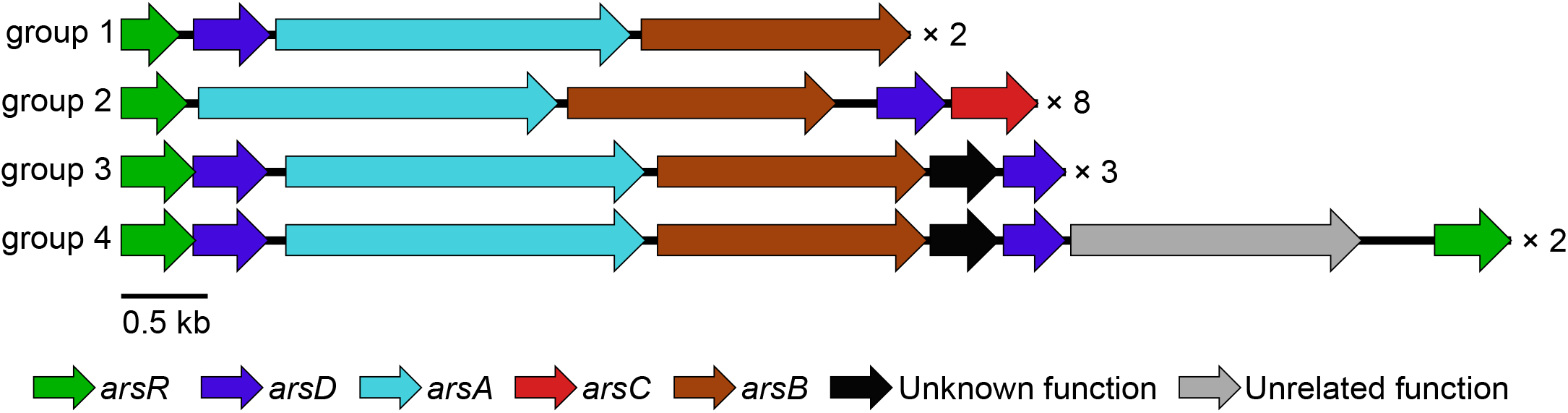
Arsenic resistance gene clusters identified in 15 *Lactobacillaceae* plasmids. Numbers on the right side of the clusters indicate the number of plasmids showing that types of arrangement of the cluster. *arsR*, a regulator; *arsA*, ATPase component; *arsB*, secondary transporter; *arsD*, transcriptional regulator; and *arsC*, arsenate reductase. The plasmids represented in each group were from, group 1: *L. sakei* WiKim0074 (CP025207.1), *L. curvatus* ZJUNIT8 (CP029967.1); group 2: *L. hokkaidonensis* LOOC 260 (AP014681.1), *L. buchneri* NRRL B-30929 (CP002653.1), *L. plantarum* CLP0611 (CP019723.1), *L. brevis* ZLB004 (-) (CP021460.1), *L. plantarum* HAC01 (-) (CP029350.1), *L. plantarum* DR7 (-) (CP031319.1), *L. plantarum* ATG-K6 (CP032465.1), *L. plantarum* ATG-K8 (-) (CP032467.1); group 3: *L. plantarum* TMW 1.277 (CP017364.1), *L. paraplantarum* DSM 10667 (CP032747.1), *L. plantarum* WCFS1 (CR377166.1); group 4: *L. plantarum* MF1298 (-) (CP013150.1 and CP013151.1). Details are available in the supplementary Table S9.

Several genes related to uptake of the potassium were found in the plasmids of *L. plantarum* and *Lactiplantibacillus pentosus* which are NL (35 genes) and *L. brevis* which is a FLL (10 genes); whereas, none of the plasmids from VAL had this gene (Table S5). This observation is consistent with the earlier report suggesting the potassium channels are not essential in the host-dependent bacteria and are found mostly in the free-living and metabolically versatile bacteria (45). Genes encoding manganese transport protein were found to be enriched in plasmids of NL and mostly had *dtxR* family transcriptional regulator on the upstream side. (Table S12). Other genes enriched on the plasmids from NL mainly include chloride channel protein, calcium-transporting ATPase, magnesium transporter, and Na+/H+ antiporter (Table S5). These proteins are involved in functions such as acid resistance, salt tolerance, metal ion homeostasis, and oxidative stress tolerance (46), (47), (48), probably suggesting their contribution to adaptation of NL to the dynamic environments.

The *dps* gene per strain was 10 and 3 fold more abundant on the plasmids from FLL (found in four species) and NL (two species), respectively, as compared to VAL (one species) (Fig. 4d). Dps is involved in tolerance towards oxidative, pH, heat, and irradiation stresses as well as metal toxicity and attachment to abiotic surfaces during the stationary phase in *E. coli* (49) and has been thought to protect *L. plantarum* GB-LP2 from the oxidative damage (50). Higher abundance of this gene in the plasmids from FLL and NL reflects the association of these strains with the open and harsh environment where exposure of such stresses is comparatively higher.

### 3.3.4 Cell wall/membrane/envelope biogenesis

Within this category, heteropolysaccharides (HePS) biosynthesis (246 CDSs), adherence (89 CDSs), and cell wall biosynthesis and degradation (77 CDSs) were the major subcategories. HePS biosynthesis was represented by 124 CDSs of glycosyltransferases (GTs), 21 CDSs of tyrosine kinase modulator (*epsB*), 53 genes of polysaccharide synthesis proteins putatively involved in HePS biosynthesis, and 48 genes involved in the biosynthesis of activated sugar precursors (Fig. 4e, Table S5). The proportion of these CDS per strain involved in HePS biosynthesis was 34 and 35 fold and significantly higher in the plasmids of NL and FLL, respectively, as compared to VAL. The significant difference between VAL and NL was also reflected in the species-level comparison (Fig, 4d, Table S6). As some of the GTs were also represented in other COG categories such as Carbohydrate transport and metabolism and Function unknown, their comprehensive identification in the *Lactobacillus* plasmids was performed using dbCAN2 meta server (Table S13). A total of 99 GTs belonging to nine families were found on 47 plasmids with 33 plasmids having two or more GTs with the dominance of GT2 family (52% of the CDSs).

We further analyzed the plasmids for the presence of the HePS biosynthesis gene clusters. Out of 512 plasmids, HePS genes clusters were found on 16 plasmids of 16 strains represented by six species (Fig. 6, Table S14). Of these, four clusters had all the genes required for HePS biosynthesis (25). Although a large number (12) of clusters analyzed in this way were incomplete, they can contribute to the HePS production in collaboration with other EPS clusters present in the chromosomes of the same strain. Such contribution of multiple HePS clusters within a strain to various properties of HePS has been demonstrated in *L. plantarum* WCFS1 (51). The absence of *epsA* in the HePS clusters from *L. plantarum* plasmids is consistent with our earlier similar observation in case of *L. plantarum* chromosomes (25). HePS is important in many VAL for colonization in the gut and in few cases withstanding low pH and high bile concentration in the stomach (52). The higher proportion of HePS related genes in the plasmids from NL and FLL than VAL probably suggests that HePS might be even more important for NL and FLL in adapting to the more diverse and harsh environments. This hypothesis is also supported by a recent genome analysis showing that the capsular exopolysaccharide encoding genes occur more frequently in the environmental bacteria than the pathogens (53).

**Figure 6:**
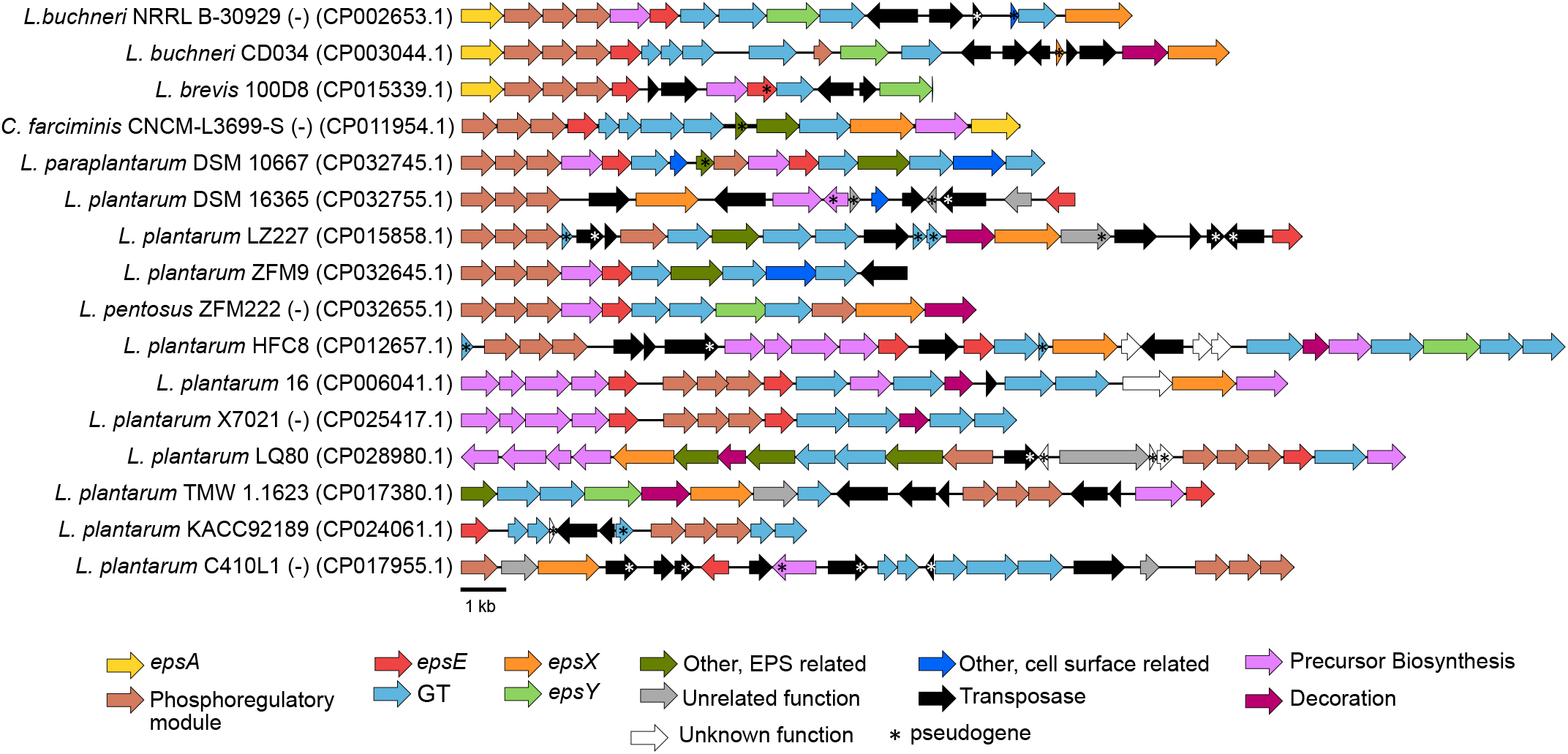
EPS gene clusters identified in 16 *Lactobacillaceae* plasmids. The clusters encoded by the negative strand are represented by the negative sign in the parentheses after the strain name. The Genbank accession number of the plasmid has been mentioned in the parentheses after the strain name.

A large number (143) of glycosyl hydrolases (GH) were also detected under in the plasmids and were classified in 19 CAZy families (Table S15) according to the dbCAN2 meta server. The contributions of GH1 (16.7%) and GH13 (15.3%) families were maximum followed by GH25 (13.9%). The proportion of the GH genes on the plasmids was significantly higher in FLL and NL as compare to VAL. Almost all the GH68 (fructansucrases) and GH70 (glucansucrase) which are involved in the production of homopolysaccharides were found only on the plasmids of *L. plantarum*. Moreover, some of the *L. plantarum* strains had multiple of such genes. For example, *L. plantarum* 16 had GH68 as well as GH70 on the same plasmid and a complete heteropolysaccharide biosynthesis gene cluster on another plasmid. Similarly, *L. plantarum* subsp*. plantarum* TS12 had two GH68, one on each of the two plasmids and also had a GH70 on another plasmid. This is consistent with the higher occurrence of the genes involved in the HePS biosynthesis in the plasmids of NL as described above. It is interesting to note the absence of glucansucrase on the plasmids VAL species but their presence in the chromosome of *Limosilactobacillus reuteri* which is a VAL (54).On the contrary, we found absence of glucansucrase in the chromosomes of *L. plantarum* 16 and *L. plantarum* subsp*. plantarum* TS12, which had these genes on the respective plasmids. This observation probably suggests the obligatory requirement of glucansucrase for the lifestyle adapted by limosilactobacilli and its conditional requirement in case of *L. plantarum*. The higher abundance of the lysozyme family (GH23 and GH25) and N-acetylglucosaminidase (GH3 and GH73) genes in the plasmids of FLL and NL might help them in hydrolyzing the cells walls of the competing bacteria in the open environment. Such a higher abundance of lysozymes encoding genes in the environmental isolates than the host-associated isolates has also been reported in *E. coli* (55). Similarly, higher metabolic versatility of NL and FLL is highlighted by the abundance in their plasmids of GH1 and GH2 which enable utilization of various oligosaccharides and GH13 and GH65 which enable utilization of starch.

We analyzed whether *Lactobacillaceae* plasmids encode any of the proteins required for adhesion to the intestinal epithelial (56). Based on annotated data and manual curation, 91 genes putatively involved in adhesion were observed in the lactobacilli plasmids. These genes mainly encoded cell wall anchor domain protein (39 CDSs), sortase (34 CDSs) and collagen-binding surface protein (16 CDSs) (Table S4). There was no significant difference between the habitats for the proportion of the plasmidome encoding these genes collectively. However, a higher number of the genes for collagen-binding surface protein were found on the plasmids of VAL (9 CDSs) as compare to NL (5 CDSs) and FLL (2 CDSs). This protein is involved in adhesion of the lactobacilli to the mammalian extracellular matrix (57). The proportion of sortase and cell-wall anchor domain proteins was higher in the plasmids from NL (37 CDSs) than VAL (5 CDSs). Sortases and sortase dependent proteins have been mostly studied for their role in interactions with the host cells. However, large numbers of these genes have already been reported in NL, such as *L. plantarum* and have been correlated to the interactions with the dynamic environments (58).

#### 3.3.5 Defense systems

The proportion of Defense systems (V) genes was significantly higher in the plasmids of VAL species as compared to NL. The major subcategories observed were restriction-modification (R-M) system (275 CDSs), toxin-antitoxin (TA) (263 CDSs), ABC transporters (123 CDSs) and CRISPR-Cas associated protein (14 CDSs) (Fig. 4f, Table S5). Of the four major types (I-IV) of R-M systems, the type I R-M system was found to be the most abundant with 175 CDSs. Although the fraction of type I R-M system genes was overrepresented on the plasmids of VAL, all the three essential subunits (R, M, S) were present in the highest numbers in FLL (9) and NL (8) as compared to the VAL (4). The average proportion of TA genes on the plasmids was significantly higher in FLL (found in seven species) and NL (found in four species) as compared to VAL (found in four species). TA loci are involved in regulating the gene expression under stress conditions and are thought to be more important to the free-living bacteria since they are more frequently under nutritional stress than the host-associated bacteria (59) justifying the above observation.

The proportion of genes encoding ABC transporters was 4.2 and 3 fold higher in the plasmids of VAL than those from FLL and NL, respectively (Fig. 4f). Most of the ABC transporters under this COG category appear to be involved in the transport of the antimicrobial peptides and antibiotics outside the cell (Table S5). Higher proportion of such genes in the plasmids of VAL is similar to the overrepresentation of the MFS transporters as discussed above and bacteriocin-encoding genes (see section antibacterial activity) in the similar plasmids. This observation also suggests the contribution of plasmids in defense against other bacteria in the more competitive environment and effluxing the antibiotics.

#### 3.3.6 Amino acid transport and metabolism

A total of 387 CDSs were found under Amino acid transport and metabolism (E) with no significant difference across the habitats. Amino acids metabolism (161 CDSs), osmoprotectant transport system (72 CDSs) and peptidase (16 CDSs) were the major subcategories (Fig. 4g, Table S5). *L. salivarius* plasmids uniquely had several genes involved in the amino acid metabolism such as 3-dehydroquinate dehydratase, serine dehydratase, and cysteine desulfurase (Table S5). The genes required for the conversion of methionine to cysteine, including cysteine synthase, cystathionine lyase, and serine acetyltransferase were found only in a few *L. paracasei, L. casei, L. rhamonosus and Limosilactobacillus fermentum* plasmids and this is consistent with this well-characterized pathway in *L. parcasei* (60).

Some plasmids encoded peptidases and their fraction was significantly higher in the plasmids from VAL as compared to NL and FLL. However, these genes were confined to the plasmids from *L. salivarius* in VAL group. A total of 68 glycine betaine transporter genes were found on the plasmids, of which 64 belonged to NL (*L. plantarum* and *L. pentosus*, Table S4). Glycine betaine transporters are involved in providing osmoprotection to the bacteria (61) suggesting that plasmids may play a role in the similar function in NL.

#### 3.3.7 Energy production and conservation

The category Energy production and conservation (C) was significantly over-represented in the plasmids of VAL as compared to NL. We further classified the genes under this category as those associated with aerobic metabolism (98 CDSs), anaerobic metabolism (71 CDSs), and others (97 CDSs) (Fig. 4h, Table S5). Plasmids from VAL had a significantly higher proportion of the genes associated with anaerobic metabolism than NL and none of FLL had such genes present on their plasmids (Fig. 4h). Although two genes, *viz*., pyruvate formate lyase and acetaldehyde dehydrogenase were present only in *L. salivarius*, other genes such as lactate dehydrogenase, alcohol dehydrogenase and fumarate reductase were each present in two or more VAL species such as *L. salivarius*, *L. amylolyticus*, *L. amylovorus*, *L. reuteri* and *Lactobacillus gasseri*. There was no significant difference between the habitats for the proportion in the plasmids of the genes involved in the aerobic metabolism. However, genes encoding the oxygen utilizing enzymes *viz*., NADH oxidase and pyruvate oxidase were present only in NL (*L. plantarum* and *L. paraplantarum*) and FLL (*Paucilactobacillus hokkaidonensis, L. brevis, S. paracollinoides, Lentilactobacillus parabuchneri*). Although *Lactobacillus* species are generally considered to be aerotolerant, because of the anaerobic gut environment, most of VAL such as *L. johnsonii* and *L. gasseri* are considered strictly anaerobic (62), (63). On the other hand, NL and FLL exhibit respiratory growth (64). Thus, NADH oxidase can help FLL and NL in the regeneration of NAD^+^ using oxygen as an electron acceptor. Furthermore, *Lc. lactis* NADH oxidase was also shown to be inactivated under anaerobic conditions or low pH (65) justifying its dispensability in VAL. Similarly, the anaerobic metabolism genes which were dominant in the plasmids from VAL can assist in the maintaining the redox balance under the anaerobic conditions in these species.

#### 3.3.8 Stress resistance

To confront a variety of harsh conditions, lactobacilli are equipped with an arsenal of stress signaling pathways (66). To get insights into the possible contribution of plasmids in the stress tolerance in *Lactobacillaceae*, we explored whether any of the various stress tolerance-associated genes described earlier in *Lactobacillus* (67), (68), (69), (70), (71) were present on the plasmids by evaluating the eggNOG, NCBI, and KEGG annotations. In total 228 stress resistance genes related to oxidative stress resistance (104 CDSs), multiple stress resistance (*clp*; 39 CDSs), universal stress resistance protein (*uspA*; 26 CDSs), stress response regulator protein (*gls24*; 15 CDSs), stress response membrane protein (GlsB/YeaQ/YmgE family 11 CDSs), low pH tolerance genes (13 CDSs) and bile tolerance (10 CDSs) were detected on 103 plasmids across 62 different strains (Table S16).

Within oxidative stress tolerance, genes encoding thioredoxin (TrxA, 33 CDSs), thioredoxin reductase (TrxB, 20 CDSs), glutathione reductase (GshR, 28 CDSs), and disulfide isomerase (FrnE, 20 CDSs) were detected in the *Lactobacillaceae* plasmids (Table S17). The functions of *gshR*, *trxA*, and *trxB* have already been established in NL and FLL (14), (72). *FrnE* encodes for a disulfide bond formation protein (DsbA) belonging to the thioredoxin family which plays a role in oxidative stress tolerance and has been characterized in *Deinococcus radiodurans* (73). Interestingly, 17 plasmids which were solely from NL and FLL had all the four genes, *trxA*, *trxB*, *gshR*, and *frnE* present in the same order and had *arsR* upstream of *trxA* (Fig. 7b, Table S18) suggesting the possible involvement of this transcriptional regulator in the expression of all the downstream genes associated with the oxidative stress response. Such a contribution of an ArsR family transcriptional regulator in the expression of *trxA2* upon the exposer to the oxidative stress has been reported in Cyanobacterium *Anabaena sp.* PCC 7120 (74). This observation along with the collective overrepresentation of these genes in the plasmids of FLL as compared to HAL indicates the contribution of these plasmids in relieving the oxidative stress in FLL and NL which is more exposed to the aerobic conditions.

**Figure 7:**
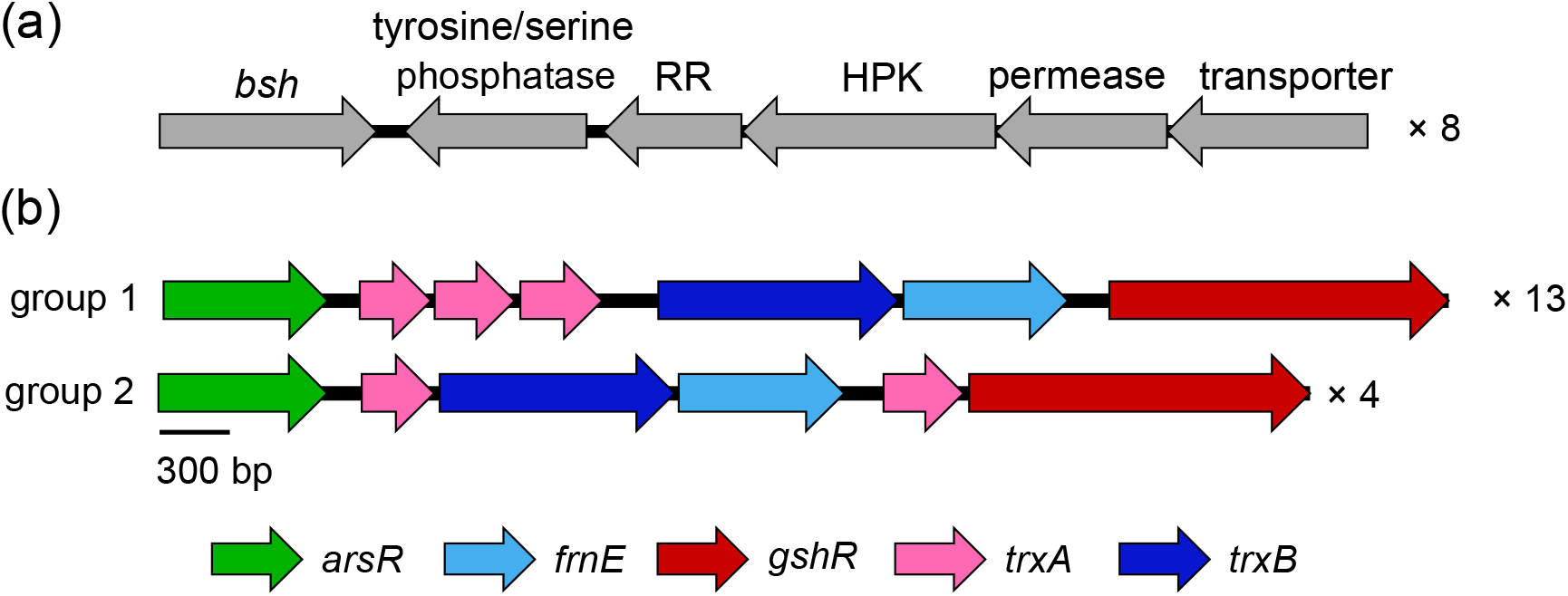
Organization of the (a) bile and (b) oxidative stress resistance genes in *Lactobacillacea*e plasmids. Numbers on the right side of the clusters indicate the number of plasmids showing that types of arrangement of the cluster. The bile resistance genes (a) were *bsh*, choloylglycine hydrolase; phosphatase, protein tyrosine/serine phosphatase; RR, two-component response regulator; HPK, signal transduction histidine kinase; permease, ABC transporter permease protein; and transporter, ABC-type multidrug transport system ATPase component. The bile resistance genes were from *L. salivarius* UCC118 (CP000234.1), *L. salivarius* CECT 5713 (CP002037.1), *L. salivarius* JCM1046 (CP007647.1), *L. salivarius* str. Ren (CP011404.1), *L. salivarius* CICC 23174 (CP017108.1), *L. salivarius* ZLS006 (CP020859.1), *L. salivarius* BCRC 12574 (CP024065.1), *L. salivarius* BCRC 14759 (CP024068.1). The oxidative stress resistance genes (b) were *arsR*, transcriptional regulator; *trxA*, thioredoxin; *trxB*, thioredoxin reductase; *frnE*, disulfide isomerase; and *gshR,* glutathione reductase. The oxidative stress resistance genes were from, group 1: *L. plantarum* LP3 (CP017068.1), *L. plantarum* CAUH2 (-) (CP015129.2), *L. brevis* LMT1-73 (CP033887.1), *L. plantarum* LMT1-48 (CP033892.1), *L. brevis* SRCM101174 (CP021481.1), *L. brevis* 100D8 (CP015340.1)*, L. brevis* KB290 (-) (AP012172.1), *L. brevis* SRCM101106 (CP021676.1), *L. plantarum* ATG-K2 (-) (CP032463.1), *L. plantarum* DSR M2 (CP022293.1), *L. sakei* WiKim0063 (CP022710.1), *L. sakei* WiKim0072 (-) (CP025138.1), *L. sakei* WiKim0074 (CP025207.1); group 2: *L. plantarum* ZFM55 (CP032360.1), *L. plantarum* ZFM9 (CP032643.1), *L. plantarum* SRCM102022 (CP021502.1), *L. plantarum* KACC 92189 (CP024060.1).

A total of 39 Clp proteinase genes (*clpL*, *clpV*, *clpX, clpC*) were detected with their fraction significantly higher in the plasmids of FLL as compared to those from HAL (Table S16). The primary function of the Clp complex is to degrade the abnormal or misfolded proteins under stressful conditions (75), (76) with its increased expression observed in *L. plantarum* WCFS1 under heat stress condition (77). The total of 52 genes encoding other stress response proteins *viz*., UspA, Gls24 *and* GlsB/YeaQ/YmgE family protein were found on the *Lactobacillaceae* plasmids of which only two ORFs were encoded by the plasmids of VAL. The gene *uspA* is considered as universal stress resistance gene and was shown to respond to various stresses such as heat shock, oxidants, DNA damage, and nutrient starvation in *L. plantarum* (78). Similarly, gene *gls24,* encodes a stress response regulator protein and is induced under copper stress in *Enterococcus hirae* (79).

*Bsh* encoding choloylglycine hydrolase involved in the deconjugation of bile salts in the mammalian gut disabling their inhibitory effect was present only in the plasmids from VAL (8 genes) and FLL (2 genes), though confined only to one species in each habitat (Table S4). Interestingly, all these plasmids also had a two-component system encoded by histidine kinase and response regulator genes, and ABC transporter genes downstream of *bsh* though encoded by the opposite strand (Fig. 7a, Table S19). An operon containing *bsh* and the two-component system genes was shown to be involved in bile salt resistance in *L. acidophilus* (80). In addition to these, an ABC transporter locus was shown to be present next to such bile resistance operon in *L. gasseri* (81). Thus, in light of these facts and few other previous studies showing strong correlation between the bile salt resistance genotype, phenotype, and the habitat of the LAB (82), it appears that plasmids play an important role in survival of at least in some lactobacilli during the passage through GI tract.

#### 3.3.9 Antibiotic resistance

Since antibiotic resistance is one of the safety concerns associated with LAB used for human consumption (83), we examined the extent of the occurrence of ARGs on the *Lactobacillaceae* plasmids using Comprehensive Antibiotic Resistance Database (CARD). A total of 58 ARGs were found on 48 plasmids in 42 *Lactobacillus* strains (Table S20). These ARGs encoded resistance for lincosamides (26 *lmrB*, two *lnuA*, and one *lsaA*), tetracycline (15), multiple antibiotics (6), aminoglycoside (2), chloramphenicol (2), erythromycin (1), pleuromutilin (2), streptogramin (1), and vancomycin (1). The presence of *lmrB* has been previously shown in *L. gasseri* UFVCC 1091 (84) and was also found to be involved in bacteriocin secretion and immunity in *Lc. lactis* (85). Campedelli et al. (86) reported the presence of 18 tetracycline resistance genes in the raw data of the genomes of 182 type strains of *Lactobacillus*. Thus, the presence of 15 tetracycline resistance genes in 512 plasmids in the current set appears to be a much higher proportion. Of these, eight genes encoded for the efflux pumps (Tet (L) and Tcr3); whereas seven encoded for ribosomal protection proteins (TetM, TetW, and TetB(P)). Majority of the tetracycline resistance genes (eight) were found on the plasmids of VAL and this phenomenon could be related to the usage of the antibiotics as tetracycline is one of the most commonly used antibiotics in humans. This hypothesis is also supported by an earlier report suggesting that the tetracycline resistance genes in the human intestinal bifidobacteria might have been acquired from the intestinal pathogens (87). Although it is important that LAB for the food and probiotic applications should be free of antibiotic resistance and the associated genes, some of the previous reports have shown the absence of correlation between the antibiotic resistance phenotype and genotype (86), (88). Thus, the mechanism of antibiotic resistance in the strains lacking the related ARGs and that of the antibiotic susceptibility in spite of having the ARGs needs to be deeply studied in *Lactobacillaceae*.

ARGs are often reported to co-occur with HMRGs in many bacteria especially under the high concentration of heavy metals in the environment (89). Thus, we analyzed the co-occurrence of ARGs with HMRGs. No significant difference was found in the proportion of plasmids having ARGs with or without HMRGs (logistic regression, p < 0.05, data not shown). Such a low extent of co-occurrence of these genes has also been reported earlier in *Lactobacillus* and *Lactococcus* (90).

#### 3.3.10 Antibacterial activity

Lactobacilli are known as one of the important producers of bacteriocins which show antimicrobial activity against closely related bacteria. To identify bacteriocin encoded ORFs in lactobacilli plasmids, the plasmid genomes were screened with the help of BAGEL4 database (91). A total of 18 plasmids of 18 strains were found to encode genes for bacteriocin operons. Twelve plasmids contained complete operons collectively for class IIa (4 plasmids), IIb (5 plasmids), and IIc (3 plasmids) bacteriocins and encoded genes for structural peptides and immunity and transport proteins (Table S21a). In total, 27 structural genes were found on the plasmids, and nine of them appear to encode novel bacteriocin (identity < 85%) (Table S21b). The highest number of the structural genes (24) encoded class II bacteriocins, whereas only one plasmid (*Lactobacillus gallinarum* HFD4) encoded three bacteriolysin genes (class III) (Table S8b). This is consistent with the earlier observation in *Lactobacillus* (92). Although the presence of structural genes alone in a bacterium is not sufficient for the production and secretion of the bacteriocins, complementation of such genes with the accessory genes from other strains was shown to support the secretion of the novel bacteriocins (93). This suggests the potential application of *Lactobacillaceae* plasmids in the production of novel bacteriocins. In the context of habitats, the proportion of strains having complete bacteriocin operons as well as those having structural genes in the plasmids was highest in VAL (18.5% and 22.2%, respectively) than NL (9.5% and 13.6%, respectively) and none of the FLL had the complete bacteriocin operons or structural genes. A similar observation was reported earlier with a high prevalence of bacteriocin encoding genes in the *Lactobacillus* strains isolated from animal and human gut than those isolated from other sources (92). This observation also suggests that plasmids might offer a competitive advantage to the lactobacilli, mostly in the host-adapted environment.

## 4. Conclusion

Plasmids have been shown to be important for the niche-adaptation in many individual bacteria. We demonstrated this phenomenon in *Lactobacillaceae* by large-scale comparative genomic analysis of the publically available plasmid sequence data. The genomic content of the plasmids was consistent with the respective lifestyle adopted by lactobacilli suggesting that the plasmids might enhance the niche-specific fitness of the strains. Whether the plasmid-encoded genes are redundant as the chromosomal ones remains to be determined. Provided that numerous bacterial genes have no proved function assigned to them, at least some of the smaller plasmids with a very few genes offer a readymade tool to understand the function of these genes. Specifically, after incorporating suitable selection marker into the plasmids and their transformation into a model LAB, the altered phenotype can give readout of the function of such genes. Additionally, some of the genes present on the plasmids such as those conferring resistance to antibiotics, heavy metals, oxidative stress, bacteriocins, etc. as well as those ascribing certain biochemical traits such as utilization of specific sugars can be developed as the selection markers.

Considering the arsenal of the genes present on the plasmids and the commercial importance of the *Lactobacillaceae* species, plasmids have huge potential in improving the properties of other strains of these bacteria. For example, the plasmids with PTS systems can enable utilization of broad range of sugars, those with the stress resistance genes can enable bacterial sustenance under industrial as well as gastrointestinal settings, those with EPS biosynthesis related genes can improve the food-related properties, those with the genes involved in redox balancing can assist in the production of value-added chemicals, and so on. Taken together, this study not only provides a bird’s eye view of the plasmidome of *Lactobacillaceae* and fundamental insights into their ecological importance but also opens new avenues to understand the biological functions and invent novel biotechnological applications of the plasmids.

## Authors and contributors

Conceptualization, R.K.; Methodology, D.D. (Dimple Davray); Formal analysis, D.D. (Dimple Davray), D.D. (Dipti Deo); Investigation, D.D. (Dimple Davray); Resources, R.K.; Writing – original draft preparation, D.D. (Dimple Davray); Writing – review and editing, R.K.

---

Figure S1: Heatmap based on the presence (black areas) and absence (white areas) of protein families encoded by the *Lactobaccilaceae* plasmids as identified by MCL analysis.

Figure S2: BLAST comparisons using EasyFig program of the highly similar plasmids identified in different strains of *Lactobacillaceae* based on the Sørensen–Dice similarity coefficient. PF numbers in the parentheses after the gene names indicate MCL protein families.

