## Supplementary material for "Plasmids encode niche-specific traits in *Lactobacillaceae*": Fig. S1

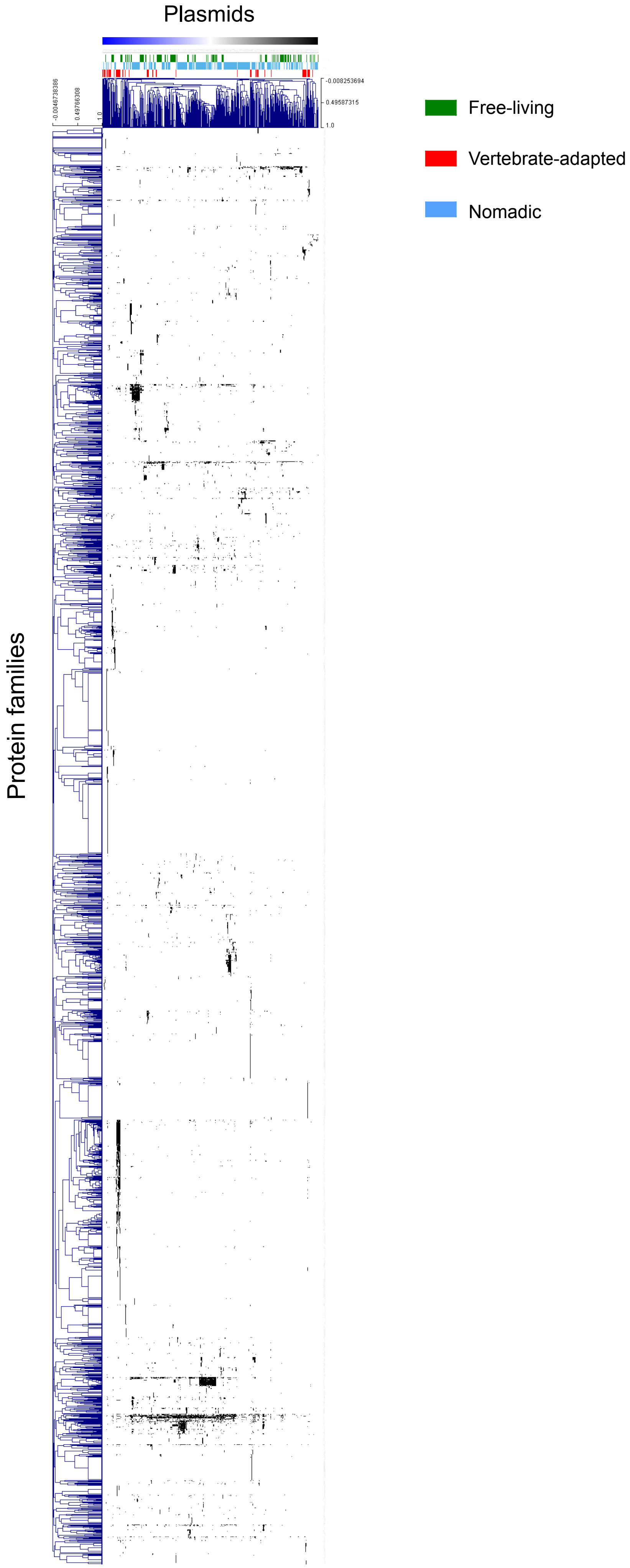


Figure S1: Heatmap based on the presence (black areas) and absence (white areas) of protein families encoded by the *Lactobaccilaceae* plasmids as identified by MCL analysis.
