## Supplementary material for "Plasmids encode niche-specific traits in *Lactobacillaceae*": Fig. S2

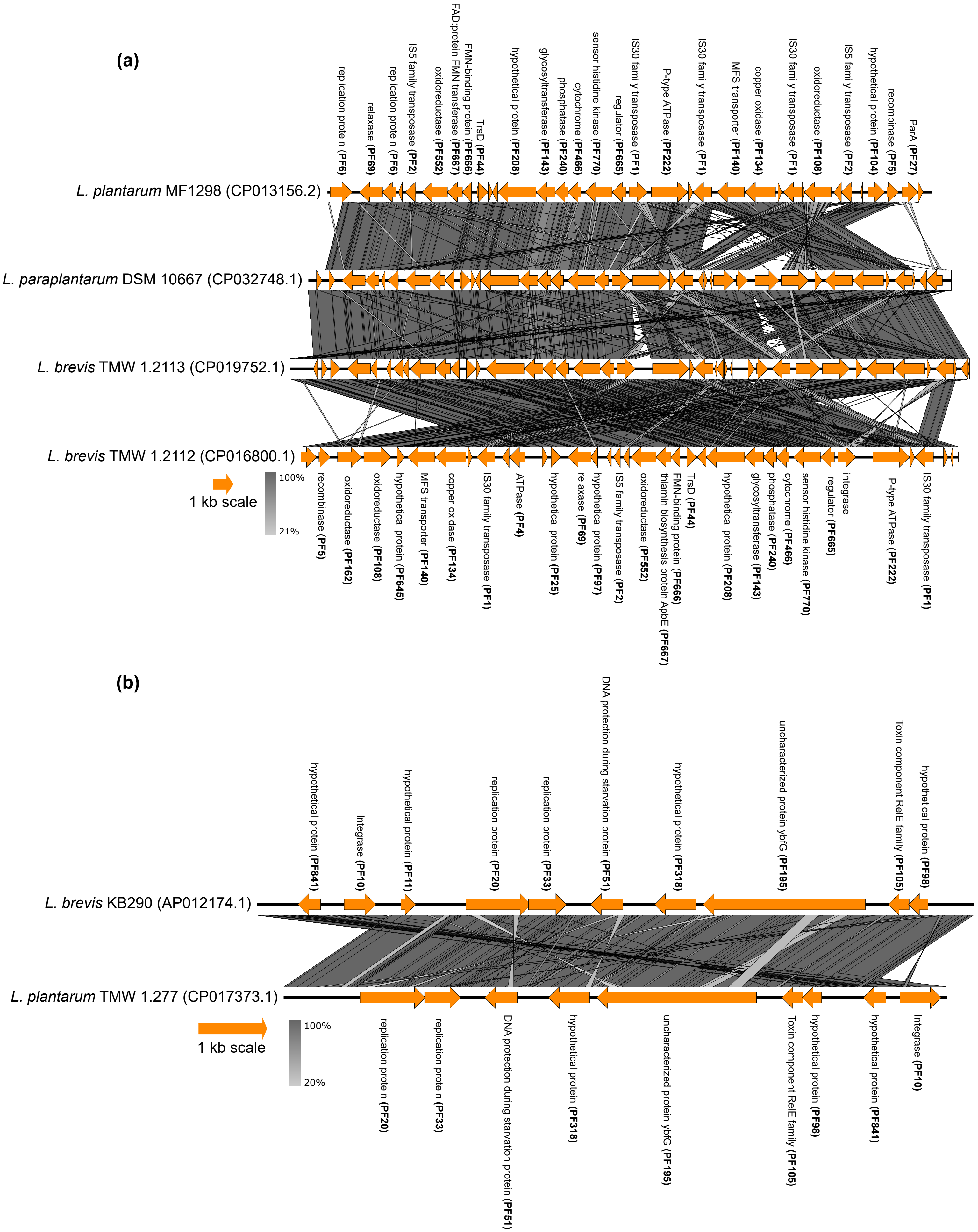

(a)
replication protein (PF6)
relaxase (PF69)
replication protein (PF6)
IS5 family transposase (PF2)
oxidoreductase (PF552)
FAD:protein FMN transferase (PF667)
FMN-binding protein (PF666)
TrsD (PF44)
hypothetical protein (PF208)
glycosyltransferase (PF143)
phosphatase (PF240)
cytochrome (PF466)
sensor histidine kinase (PF770)
regulator (PF665)
IS30 family transposase (PF1)
P-type ATPase (PF222)
IS30 family transposase (PF1)
MFS transporter (PF140)
copper oxidase (PF134)
IS30 family transposase (PF1)
oxidoreductase (PF108)
IS5 family transposase (PF2)
hypothetical protein (PF104)
recombinase (PF5)
ParA (PF27)
L. plantarum MF1298 (CP013156.2)
L. paraplantarum DSM 10667 (CP032748.1)
L. brevis TMW 1.2113 (CP019752.1)
L. brevis TMW 1.2112 (CP016800.1)
1 kb scale
100%
21%
recombinase (PF5)
oxidoreductase (PF162)
oxidoreductase (PF108)
hypothetical protein (PF645)
MFS transporter (PF140)
copper oxidase (PF134)
IS30 family transposase (PF1)
ATPase (PF4)
hypothetical protein (PF25)
relaxase (PF69)
hypothetical protein (PF97)
IS5 family transposase (PF2)
oxidoreductase (PF552)
thiamin biosynthesis protein ApbE (PF667)
FMN-binding protein (PF666)
TrsD (PF44)
hypothetical protein (PF208)
glycosyltransferase (PF143)
phosphatase (PF240)
cytochrome (PF466)
sensor histidine kinase (PF770)
regulator (PF665)
integrase
P-type ATPase (PF222)
IS30 family transposase (PF1)
(b)
hypothetical protein (PF841)
Integrase (PF10)
hypothetical protein (PF11)
replication protein (PF20)
replication protein (PF33)
DNA protection during starvation protein (PF51)
hypothetical protein (PF318)
uncharacterized protein ybfG (PF195)
Toxin component RelE family (PF105)
hypothetical protein (PF98)
L. brevis KB290 (AP012174.1)
L. plantarum TMW 1.277 (CP017373.1)
1 kb scale
100%
20%
replication protein (PF20)
replication protein (PF33)
DNA protection during starvation protein (PF51)
hypothetical protein (PF318)
uncharacterized protein ybfG (PF195)
Toxin component RelE family (PF105)
hypothetical protein (PF98)
hypothetical protein (PF841)
Integrase (PF10)

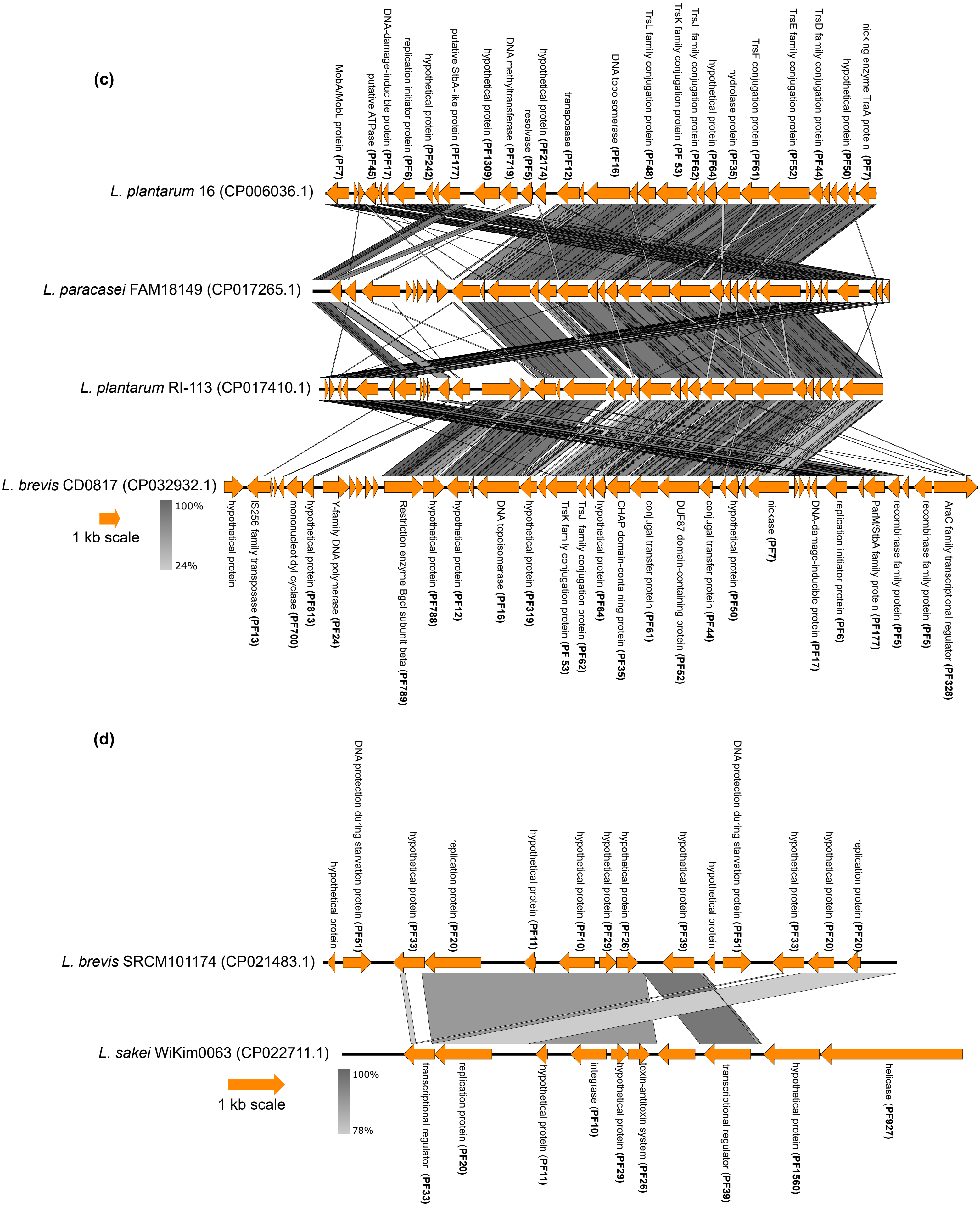

(c)
MobA/MobL protein (PF7)
putative ATPase (PF45)
DNA-damage-inducible protein (PF17)
replication initiator protein (PF6)
hypothetical protein (PF242)
putative StbA-like protein (PF177)
hypothetical protein (PF1309)
DNA methyltransferase (PF719)
resolvase (PF5)
hypothetical protein (PF2174)
transposase (PF12)
DNA topoisomerase (PF16)
TrsL family conjugation protein (PF48)
TrsK family conjugation protein (PF 53)
TrsJ family conjugation protein (PF62)
hypothetical protein (PF64)
hydrolase protein (PF35)
TrsF conjugation protein (PF61)
TrsE family conjugation protein (PF52)
TrsD family conjugation protein (PF44)
hypothetical protein (PF50)
nicking enzyme TraA protein (PF7)
L. plantarum 16 (CP006036.1)
L. paracasei FAM18149 (CP017265.1)
L. plantarum RI-113 (CP017410.1)
L. brevis CD0817 (CP032932.1)
1 kb scale
100%
24%
hypothetical protein
IS256 family transposase (PF13)
mononucleotidyl cyclase (PF700)
hypothetical protein (PF813)
Y-family DNA polymerase (PF24)
Restriction enzyme BgcI subunit beta (PF789)
hypothetical protein (PF788)
hypothetical protein (PF12)
DNA topoisomerase (PF16)
hypothetical protein (PF319)
TrsK family conjugation protein (PF 53)
TrsJ family conjugation protein (PF62)
hypothetical protein (PF64)
CHAP domain-containing protein (PF35)
conjugal transfer protein (PF61)
DUF87 domain-containing protein (PF52)
conjugal transfer protein (PF44)
hypothetical protein (PF50)
nickase (PF7)
DNA-damage-inducible protein (PF17)
replication initiator protein (PF6)
ParM/StbA family protein (PF177)
recombinase family protein (PF5)
recombinase family protein (PF5)
AraC family transcriptional regulator (PF328)
(d)
hypothetical protein
DNA protection during starvation protein (PF51)
hypothetical protein (PF33)
replication protein (PF20)
hypothetical protein (PF11)
hypothetical protein (PF10)
hypothetical protein (PF29)
hypothetical protein (PF26)
hypothetical protein (PF39)
hypothetical protein
DNA protection during starvation protein (PF51)
hypothetical protein (PF33)
hypothetical protein (PF20)
replication protein (PF20)
L. brevis SRCM101174 (CP021483.1)
L. sakei WiKim0063 (CP022711.1)
1 kb scale
100%
78%
transcriptional regulator (PF33)
replication protein (PF20)
hypothetical protein (PF11)
integrase (PF10)
hypothetical protein (PF29)
toxin-antitoxin system (PF26)
transcriptional regulator (PF39)
hypothetical protein (PF1560)
helicase (PF927)

**(e)**

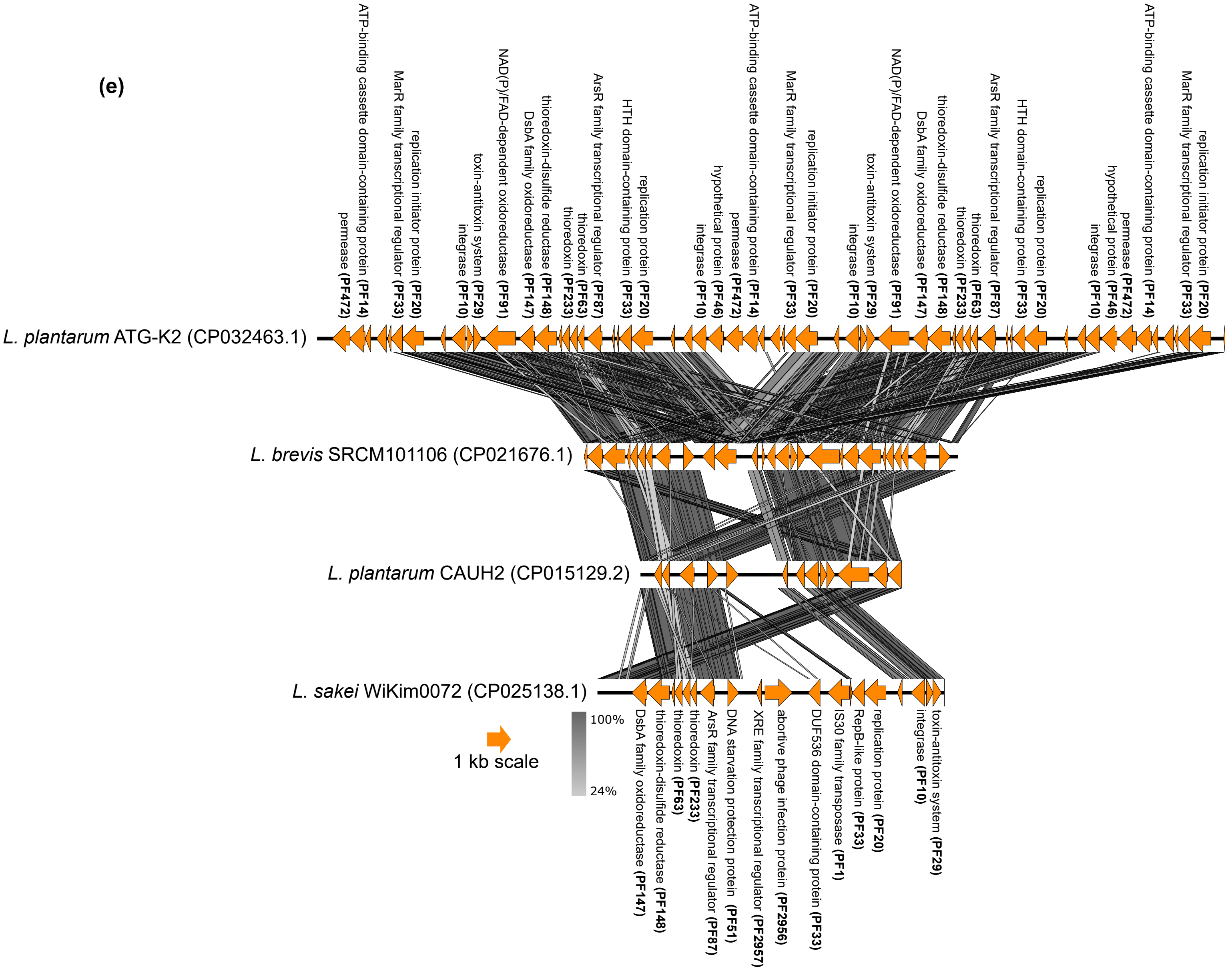


Figure S2: BLAST comparisons using EasyFig program of the highly similar plasmids identified in different strains of *Lactobacillaceae* based on the Sørensen–Dice similarity coefficient. PF numbers in the parentheses after the gene names indicate MCL protein families.
